# Loss of SynDIG4/PRRT1 alters distribution of AMPA receptors in Rab4- and Rab11-positive endosomes and impairs basal AMPA receptor recycling

**DOI:** 10.1101/2024.12.29.630674

**Authors:** Chun-Wei He, Elva Díaz

**Affiliations:** Department of Pharmacology, School of Medicine, University of California, Davis, Davis, CA, 95616, USA

**Author notes:** Correspondence: Elva Díaz.

**Keywords:** SynDIG4, PRRT1, AMPA receptor, endosomes, recycling, Rab4, Rab11

## Abstract

The transmembrane protein Synapse Differentiation Induced Gene 4 (SynDIG4) functions as an auxiliary factor of AMPA receptors (AMPARs) and plays a critical role in excitatory synapse plasticity as well as hippocampal-dependent learning and memory. Mice lacking SynDIG4 have reduced surface expression of GluA1 and GluA2 and are impaired in single tetanus-induced long-term potentiation and NMDA receptor (NMDAR)-dependent long-term depression. These findings suggest that SynDIG4 may play an important role in regulating AMPAR distribution through intracellular trafficking mechanisms; however, the precise roles by which SynDIG4 governs AMPAR distribution remain unclear. In this study, we characterized the endocytosis and recycling of GluA1-containing AMPARs under basal conditions. We did not observe any change in baseline endocytosis; however, we did observe a significant decrease in recycling of GluA1-containing AMPARs in cultured hippocampal neurons from mice lacking SynDIG4. This resulted in a significant increase in the levels of internal GluA1 and GluA2, along with greater colocalization of these subunits with Rab4-positive recycling endosomes in hippocampal neurons lacking SynDIG4. Notably, the overlap between Rab4- and Rab11-positive vesicles was elevated in hippocampal neurons lacking SynDIG4, suggesting an impairment in the trafficking between Rab4 and Rab11 compartments. Furthermore, our findings revealed a reduction in surface GluA1 within synaptic regions of hippocampal neurons lacking SynDIG4. Collectively, these results indicate that SynDIG4 regulates the distribution of GluA1-containing AMPARs via the Rab4-dependent endosomal recycling pathway, thereby maintaining AMPAR levels at synaptic regions under baseline conditions. This regulatory function of SynDIG4 may contribute to the deficits in GluA1-dependent synaptic plasticity and impairment of hippocampal-dependent learning and memory behaviors observed in SynDIG4 deficient mice.

## INTRODUCTION

The spatial and temporal organization of neurotransmitter receptors at the surface of postsynaptic sites is fundamental to efficient synaptic function, which governs how the brain processes and responds to information (1–3). At excitatory synapses, α-amino-3-hydroxy-5-methyl-4-isoxazolepropionic acid (AMPA) receptors (AMPARs) are critical for rapid synaptic transmission. Changes in the number of synaptic AMPARs reflect changes in synaptic strength under both basal and activity-dependent conditions, making them a key molecular mechanism underlying learning and memory (4–6). Thus, the regulatory mechanisms governing AMPAR trafficking within synapses is an area of intense and on-going investigation.

Many studies highlight the importance of dynamic AMPAR trafficking to and from synapses in regulating synaptic expression and modulating synaptic strength. For example, AMPARs can undergo differential sorting in endosomal compartments, rapid recycling to the plasma membrane, and lateral diffusion between synaptic and extrasynaptic sites (6–10). The endocytic machinery localized at the postsynaptic region is essential for maintaining basal synaptic transmission and plasticity to ensure the availability of a cycling pool of AMPARs near the postsynaptic density (11–13). For instance, the physical coupling of dynamin-3 to Homer anchors the endocytic zone, sustaining mobile surface AMPARs that are essential for regulating synaptic strength (14,15).

Small GTPases of the Rab family have emerged as regulators of AMPAR vesicular trafficking (16). For instance, Rab4 and Rab11 mediate the recycling of endocytosed AMPARs from sorting and recycling endosomes, respectively, during basal conditions (17,18). Impaired fusion between Rab4- and Rab11-positive endosomal compartments has been shown to reduce the surface expression of GluA1- and GluA2-containing AMPARs (19,20). These studies demonstrate that dynamic trafficking of AMPARs is essential for maintaining synaptic function and plasticity, relying on coordinated processes of endocytosis and exocytosis, via diverse endosomal sorting mechanisms to regulate their synaptic expression.

AMPAR auxiliary proteins play critical roles in regulating AMPAR distribution and gating properties (21–25). One such AMPAR auxiliary factor is Synapse Differentiation-Induced Gene 4 (SynDIG4), also known as Proline-Rich Transmembrane Protein 1 (PRRT1) (26–28), shown to be present in native AMPAR complexes through cryo-electron microscopy (EM) structural analysis (29). Notably, the transmembrane domain of SynDIG4 was observed to align approximately parallel to the transmembrane helices of the AMPAR, with the putative interaction site being the M4 helix of GluA1 (29). Consistent with these findings, the membrane-associated region of SynDIG4 has been shown to be crucial for the clustering of GluA1 and GluA2 in heterologous cells (30). Moreover, using heterologous expression in *Xenopus* oocytes, SynDIG4 was shown to modulate AMPAR gating properties in a subunit-dependent manner (31), consistent with a direct interaction. In SynDIG4 knockout (KO) mice, while total levels of AMPARs are unchanged, surface expression of GluA1 and GluA2 was significantly decreased (32). SynDIG4 overlaps primarily with extrasynaptic AMPARs (33) and localizes to early and recycling endosomes (34). Indeed, there is a significant decrease in extrasynaptic GluA1 and GluA2 in SynDIG4 KO neurons (31). Furthermore, NMDA receptor (NMDAR)-dependent long-term depression (LTD) (32) and single tetanus-induced long-term potentiation (LTP) (31) are impaired in SynDIG4 KO neurons while baseline synaptic transmission is intact. As these synaptic plasticity mechanisms require calcium-permeable AMPARs (35), which are mostly GluA1 homomers, these results suggest that SynDIG4 is required for establishing pools of extrasynaptic AMPARs (both GluA1/GluA2 heteromers and GluA1 homomers) at baseline.

These results motivated us to investigate the mechanisms by which SynDIG4 regulates the trafficking and surface expression of AMAPRs under basal conditions. We analyzed endocytosis and recycling of GluA1-containing AMPARs in wild type (WT) and SynDIG4 KO neurons and brain tissue. While basal endocytosis was unchanged, AMPAR recycling was significantly reduced in SynDIG4 KO neurons, leading to increased intracellular accumulation of GluA1 and GluA2 in Rab4-positive recycling endosomes. Furthermore, there was an increase in the overlap between Rab4-positive and Rab11-positive compartments in SynDIG4 KO neurons, suggesting impaired trafficking between them. Based on these results, we propose a model in which SynDIG4 plays a critical role in the recycling process of GluA1-containing AMPARs, particularly at the interface between Rab4- and Rab11-mediated pathways, thereby influencing their surface expression and synaptic distribution.

## MATERIALS AND METHODS

### Animals

All animal experiments were conducted in protocols approved by the Institutional Animal Care and Use Committee (IACUC) at the University of California, Davis, following the guidelines of the US Public Health Service and NIH. SynDIG4 KO mice were generated and maintained on a C57BL/6 background as described (31). Littermates of both sexes from heterozygous breeding pairs were used for experiments.

### Primary hippocampal neuron culture

Hippocampal neurons were isolated from P0∼P2 mice of both sexes and plated onto poly-L-lysine-coated glass coverslips in 6-well plates at a density of 100,000 cells per well. The neurons were cultured in astrocyte-conditioned neuron maintenance medium (NMM), consisting of Neurobasal medium (Gibco, Cat# 21103049) supplemented with GlutaMAX (Gibco, Cat# 35050079) and B27 (Gibco, Cat# A3582801). Half of the NMM volume was changed every 5 days, and neurons were fixed for experiments between days in vitro (DIV) 18 and 22.

### Bis-sulfosuccinimidyl suberate (BS^3^) surface-protein cross-linking assay

Acute coronal brain slices (400 μm thick) were obtained from P14 mice, targeting the hippocampal region, in ice-cold dissection buffer containing (in mM): 127 NaCl, 1.9 KCl, 1.2 KH_2_PO_4_, 26 NaHCO_3_, 10 D-glucose, 23 MgSO_4_, and 1.1 CaCl_2_, saturated with 95% O_2_ and 5% CO_2_. Following dissection, the slices were maintained in artificial cerebrospinal fluid (ACSF) composed of (in mM): 127 NaCl, 26 NaHCO_3_, 1.2 KH_2_PO_4_, 1.9 KCl, 2.2 CaCl_2_, 1 MgSO_4_, and 10 D-glucose, oxygenated with 95% O_2_ and 5% CO_2_ for 2 hours at 30°C to allow recovery. After recovery, the slices were transferred to 1 mL of ice-cold ACSF and treated with 40 μL of 52 mM BS^3^ at 4°C for 30 minutes with gentle mixing. The cross-linking reaction was terminated by adding 100 μL of 1 M glycine and incubating for 10 minutes. The samples were then centrifuged at 20,000 × g for 2 minutes at 4°C, after which the supernatant was removed, and the pellet was lysed for protein extraction.

### Immunoblotting

Protein samples were heated at 70°C for 10 minutes and separated by gel electrophoresis using the Bio-Rad Mini-Protean system with 4–20% gradient gels (Bio-Rad, Cat# 456-1094). After electrophoresis, proteins were transferred to nitrocellulose membranes, which were blocked in 5% milk prepared in Tris-buffered saline (TBS) with 0.1% Tween-20 (TBST) for 60 minutes at room temperature (RT). Membranes were incubated overnight at 4°C with the following primary antibodies diluted in 5% TBST: GluA1-C (Millipore, Cat# AB1504, 1:1000), GluA2-C (NeuroMab, Cat# SKU: 75-002, 1:1000), and β-tubulin (Millipore, Cat# 05-661, 1:5000). The next day, membranes were washed three times with TBST and incubated for 2 hours at RT with secondary antibodies: Goat anti-Mouse IgG1 488 (Jackson ImmunoResearch Labs, Cat# 112-545-205, 1:500), Goat anti-Mouse IgG IR700 (AzureSpectra, Cat# AC2129, 1:1000), and Goat anti-Rabbit IgG IR800 (AzureSpectra, Cat# AC2134, 1:1000). After incubation, membranes were washed three times with TBST and once with TBS at RT before imaging. Fluorescent immunoblots were imaged using the Sapphire Bioimager (Azure Biosystems, Model# Sapphire RGBNIR) and quantified with Azure Spot software (version 2.0). For analysis, the intensity of GluA1 and GluA2 bands at their predicted molecular weight region was normalized to β-tubulin. The internal-to-total ratios of GluA1 and GluA2 were compared between WT and SynDIG4 KO samples.

### Immunofluorescence

Hippocampal neurons at DIV18∼22 were fixed with 4% paraformaldehyde and 4% sucrose in phosphate buffered saline (PBS) for 7 minutes (surface labeling) or 10 minutes (standard labeling) at RT. After fixation, coverslips were washed once with ice-cold PBS and blocked with 5% fetal bovine serum (FBS) in PBS at RT for 2 hours. For surface labeling, neurons were incubated overnight at 4°C with primary antibodies before permeabilization. The antibodies used were GluA1-N (NeuroMab, SKU: 75-327, 1:100) and GluA2-N (Millipore, Cat# MAB397, 1:100). For standard labeling, neurons were permeabilized with 0.25% Triton X-100 in PBS for 8 minutes after fixation. After permeabilization, neurons were incubated at RT for 2 hours with primary antibodies. The primary antibodies included MAP2 (Millipore, Cat# AB5622-1, 1:300), PSD95 (Synaptic Systems, Cat# 124014, 1:300), GluA1-C (Millipore, Cat# AB1504, 1:200), GluA2-C (NeuroMab, SKU: 75-002, 1:200), EEA1 (Cell Signaling, Cat# 2411, 1:100), Rab4 (Cell Signaling, Cat# 2167, 1:50), Rab4 (Santa Cruz, Cat# sc-517263, 1:50), and Rab11 (Cell Signaling, Cat# 5589, 1:50), and Rab11 (Thermofisher, Cat# MA5-49197, 1:50).

After the primary antibody incubation, coverslips were washed with 0.01% Triton X-100 in PBS and incubated at RT for 2 hours with secondary antibodies. The secondary antibodies included Goat anti-Rabbit 405 (Jackson ImmunoResearch, Cat# 111-475-003, 1:100), Donkey anti-Rabbit 488 (Jackson ImmunoResearch, Cat# 711-545-152, 1:200), Goat anti-Mouse IgG2a 488 (Jackson ImmunoResearch, Cat# 115-546-206, 1:300), Goat anti-Mouse IgG2b 488 (Jackson ImmunoResearch, Cat# 115-545-207, 1:300), Goat anti-Mouse IgG2a 647 (Jackson ImmunoResearch, Cat# 115-605-206, 1:300), Goat anti-Guinea Pig 647 (Invitrogen, Cat# A-21450, 1:200), and Goat anti-Mouse IgG1 555 (Invitrogen, Cat# A-21127, 1:300). As a cell tracer, neurons were incubated with DiA (4-(4-Dihexadecylaminostyryl)-N-methylpyridinium iodide), a lipophilic dye that inserts into the plasma membranes, overnight at 4°C, where indicated.

### Antibody-feeding assay

The antibody-feeding assay was based on published protocols (9,20,36). Hippocampal neurons at DIV18∼22 were incubated with primary antibodies (GluA1-N, 1:50; GluA2-N, 1:50) diluted in conditioned media for 15 minutes at RT. Following three washes with conditioned media, the neurons were returned to a 37°C incubator for 30 minutes. For **AMPAR endocytosis experiments**, neurons were fixed and incubated with fragment antigen-binding (Fab) antibodies (80 μg/mL, Jackson ImmunoResearch Labs, Cat# 715-007-003) for 20 minutes at RT, followed by permeabilization and subsequent staining using the standard labeling protocol described previously. For **AMPAR recycling experiments**, after a 30 minute internalization step, neurons were treated with Fab antibodies (80 μg/mL) at RT for 20 minutes and washed three times with conditioned media. The neurons were then treated with dynasore (80 μM; Millipore, SML0340-5MG), a dynamin-dependent endocytosis inhibitor (37), and incubated at 37°C for 45 minutes to allow recycling of internalized, live-labeled AMPARs while inhibiting the endocytosis of the remaining surface AMPAR pool. After 45 minutes, neurons were fixed with 4% paraformaldehyde and 4% sucrose in PBS for 7 minutes, permeabilized, and subjected to the standard labeling protocol.

### Imaging and analysis

Images were acquired using a Leica SP8 instrument in either confocal or Stimulated Emission Depletion (STED) mode, employing a 63x or 100x objective, respectively. Z-stack images were captured throughout the entire cell volume. Prior to analysis, images were thresholded, with thresholds for each experiment determined by averaging the thresholds of at least 25% of the images within the dataset. Following maximum projection of the z-stacks, image analysis was conducted using FIJI/ImageJ (ver. 2.14.0/1.54f). For STED images, deconvolution was performed using Huygens Software (Scientific Volume Imaging) prior to thresholding. Overlap between signals was calculated using Manders Correlation Coefficient analysis via the JaCoP plugin. For quantification of the amount of surface GluA1 (sGluA1) overlapped with PSD95 puncta, colocalization of sGluA1 and PSD95 was identified by using a PSD95 mask overlaid on sGluA1 signals. The level of sGluA1 at PSD95 was quantified as the ratio of the number of sGluA1 PSD95 colocalized puncta to the total number of sGluA1 puncta. To calculate the association between sGluA1 and PSD95, cross-correlation confidence analysis in Fuji/ImageJ was performed. All the sGluA1 signals excluding the signals overlapped with MAP2 were subjected to calculation.

### Statistics

Statistical analyses were conducted using GraphPad Prism (version 10.3.1).

## RESULTS

### Intracellular pools of GluA1 and GluA2 are increased in the absence of SynDIG4

Previous studies demonstrated that surface GluA1 (sGluA1) and surface GluA2 (sGluA2) are decreased in brain lysates from SynDIG4 KO mice through surface-biotinylation assays (32). To investigate whether intracellular GluA1 (inGluA1) and intracellular GluA2 (inGluA2) are subsequently affected in SynDIG4 KO mice, we applied bis-sulfosuccinimidyl suberate (BS^3^), a membrane-impermeable reagent that covalently cross-links proteins at the plasma membrane (38), to acute brain slices obtained from WT and SynDIG4 KO mice. This method allowed the separation of surface and intracellular protein pools, which were subsequently visualized via immunoblotting. After BS^3^ treatment on one hemisphere of the brain slice, no aggregation of β-tubulin was observed, confirming that BS^3^ selectively cross-links surface proteins (**Figure 1A**). Concurrently, sGluA1 and sGluA2 formed aggregates observable in higher molecular weight regions, while inGluA1 and inGluA2 appeared at their predicted apparent molecular weight (**Figure 1A**). By comparing the ratio of the intracellular signal from the treated hemisphere to total signal intensity from the untreated hemisphere of the same slice, we found that inGluA1 and inGluA2 levels were both increased in SynDIG4 KO brain slices (**Figure 1B**).

**Figure 1.**
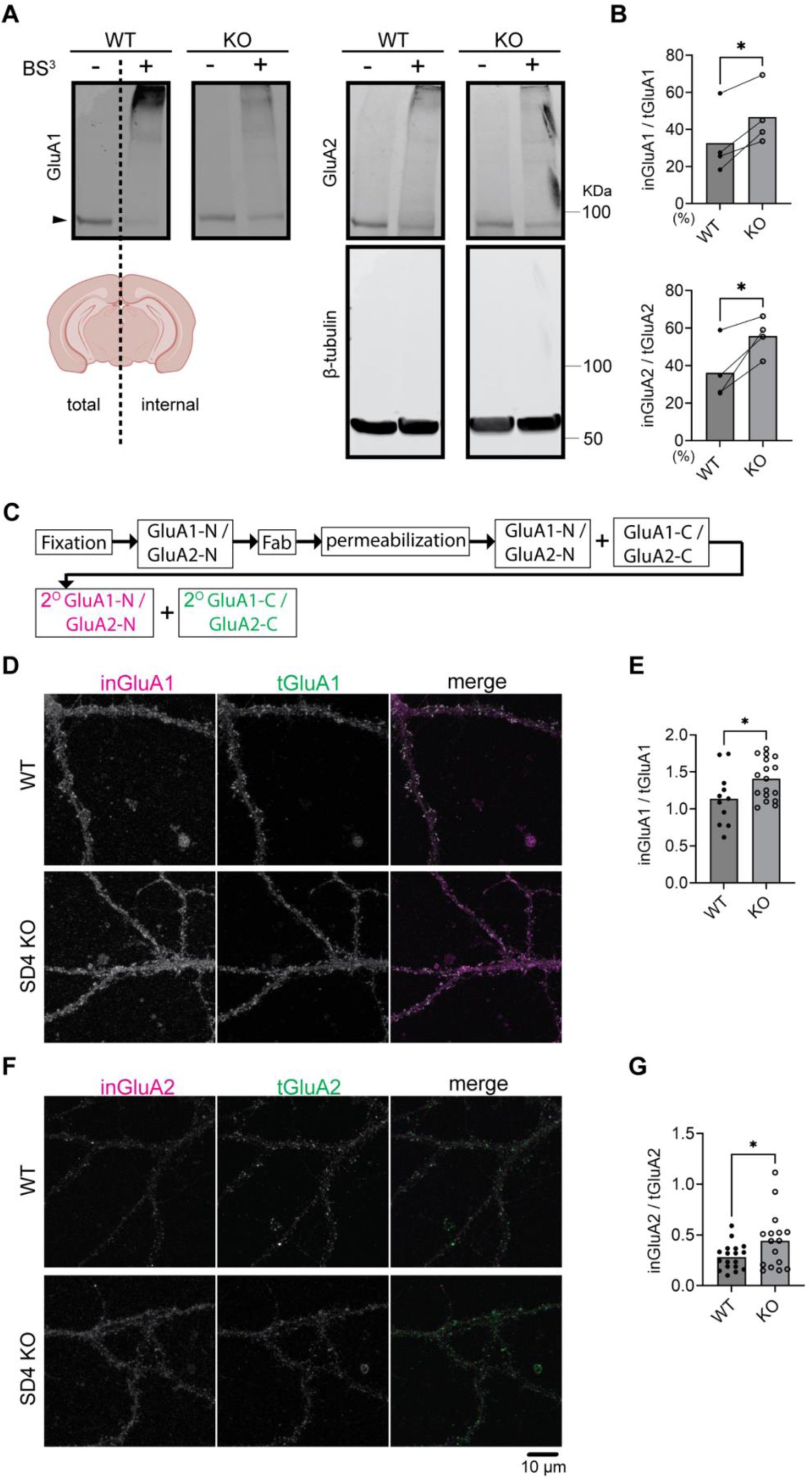
Increased Intracellular GluA1 and GluA2 Levels in SynDIG4 KO Neurons. (A) Representative immunoblots showing the separation of surface and intracellular GluA1 and GluA2 in brain slices from P14 WT and SynDIG4 KO mice after BS^3^ treatment. Arrowheads indicate the protein bands subjected to quantification. (B) Quantification of the ratio of intracellular to total GluA1 and GluA2 signals in WT and SynDIG4 KO brain slices. Data are presented as mean ± SEM, with statistical significance determined using ratio paired t-test; n = 4; *p<0.05. (C) Schematic diagram illustrating the experimental procedure for intracellular labeling of AMPARs. (D-G) Representative confocal images of WT and SynDIG4 KO neurons showing internal and total GluA1 (D) and GluA2 (F) signals. Quantification of the ratio of internal to total GluA1 (E) and GluA2 (G) signals in WT and SynDIG4 KO neurons. Data are presented as mean ± SEM, with statistical significance determined using unpaired t-test; n = 15-20; *p < 0.05. Scale bar, 10 µm.

To determine whether levels of inGluA1 and inGluA2 between acute brain slices and cultured hippocampal neurons from SynDIG4 KO and WT mice are comparable, fragment antigen-binding (Fab) antibodies were applied to fixed neurons to block surface signals by preventing secondary antibodies from accessing primary antibodies bound to the extracellular N-terminal (NT) regions of GluA1 (GluA1-N) and GluA2 (GluA2-N) prior to permeabilization (**Figure S1A**). As a control, sGluA1 and sGluA2 signals cannot be detected after adding Fab (**Figure S1B**). After permeabilization, the same primary antibodies against GluA1-N and GluA2-N followed by fluorophore-conjugated secondary antibodies were added to detect inGluA1 and inGluA2 (**Figure 1C**). By comparing the ratio of intracellular signals detected by the antibody against the NT to total signals detected by the antibody against the C-terminal (CT) regions of GluA1 (GluA1-C) and GluA2 (GluA2-C), we found that inGluA1 and inGluA2 levels were increased in cultured hippocampal neurons (**Figure 1D-G**), consistent with BS^3^ cross-linking results from brain slices (**Figure 1A-C**) and surface biotinylation results from brain lysates (32).

### SynDIG4 regulates the recycling of GluA1-containing AMPARs

Previous findings showed no significant changes in total AMPAR levels in SynDIG4 KO mice (31,32). Combined with the current results, these findings indicate that SynDIG4 likely regulates the trafficking and distribution of AMPARs rather than their expression. To pinpoint the trafficking mechanism impaired in SynDIG4 KO neurons, we employed an antibody-feeding approach to examine the endocytosis and recycling of GluA1-containing AMPARs (9,20,36). To label endocytosed GluA1 (enGluA1), live hippocampal neurons were incubated with an antibody targeting the extracellular NT region of GluA1 (GluA1-N), followed by a 37°C incubation to allow internalization of labeled receptors. Fab antibodies were subsequently applied to block residual GluA1-N epitopes, preventing further labeling. After permeabilization, antibodies against the CT region of GluA1 (GluA1-C) were applied, followed by secondary antibody incubation (**Figure 2A**). During the GluA1-N antibody-feeding process, some receptors were internalized (**Figure S2A, S2B**). After 30 minutes of internalization, the levels of enGluA1 were comparable between WT and SynDIG4 KO neurons (**Figure 2B, 2C**). To investigate recycling of GluA1-containing AMPARs, hippocampal neurons were treated with dynasore, a dynamin-dependent endocytosis inhibitor (37), following GluA1-N antibody feeding and internalization (**Figure 2D**). This treatment prevented further endocytosis, allowing recycled GluA1 (reGluA1) to reinsert into the plasma membrane. Dynasore treatment decreased enGluA1 levels in WT neurons as expected (**Figure S2C, S2D**). Interestingly, reGluA1 levels were reduced in SynDIG4 KO neurons (**Figure 2E, 2F**), indicating that SynDIG4 is important for the recycling but not for the endocytosis process of GluA1-containing AMPARs at the plasma membrane.

**Figure 2.**
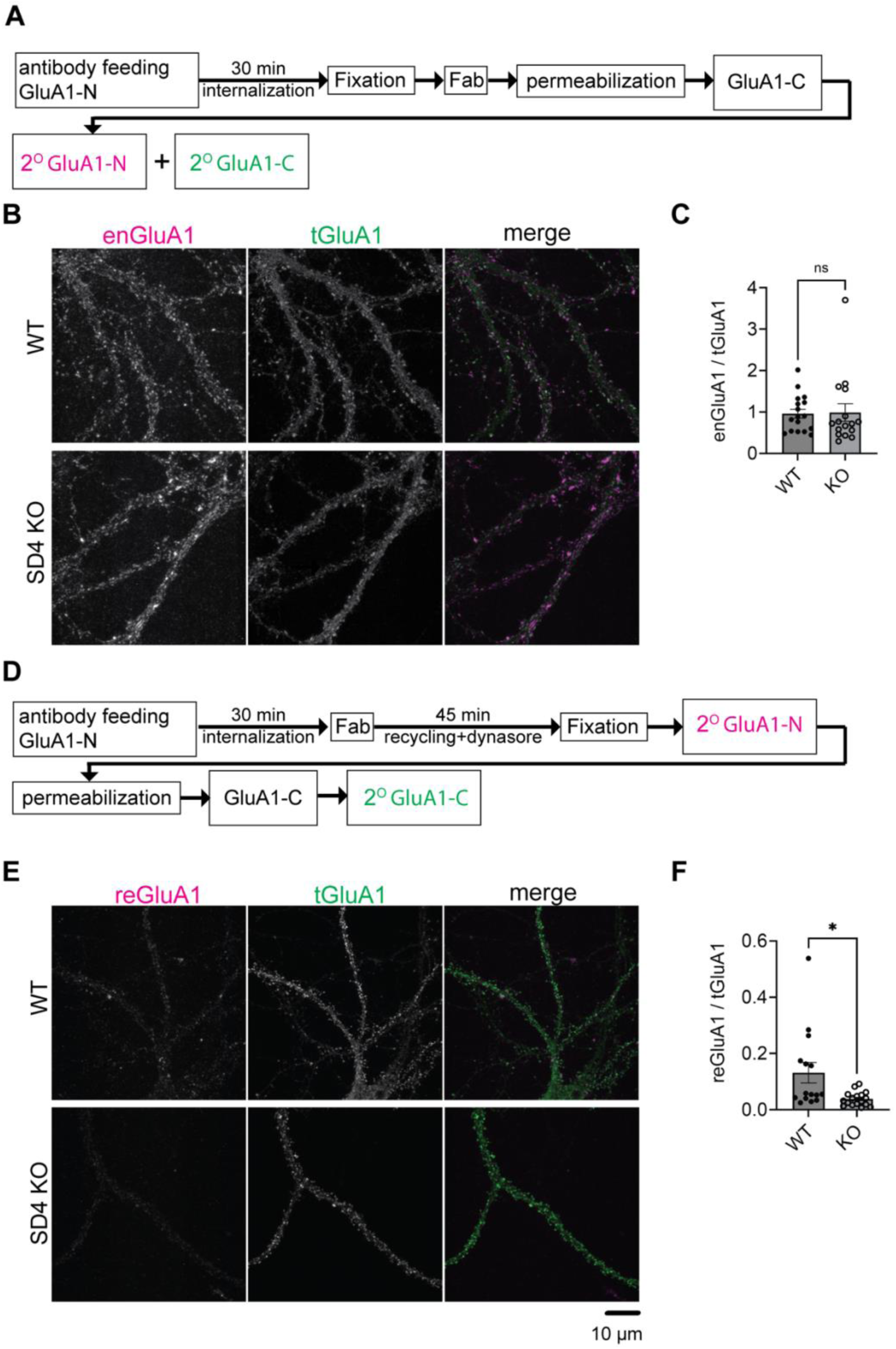
Impaired recycling of GluA1-containing AMPARs in SynDIG4 KO neurons. (A) Schematic diagram illustrating the experimental procedure for AMPAR endocytosis. After antibody feeding, neurons were incubated for 30 minutes, fixed, treated with Fab, permeabilized, and subjected to standard labeling. (B) Representative confocal images of WT and SynDIG4 KO neurons showing endocytosed and total GluA1 signals. (C) Quantification of the ratio of integrated density for endocytosed GluA1 to total GluA1 signals in WT and SynDIG4 KO neurons. (D) Schematic diagram illustrating the experimental procedure for AMPAR recycling. (E) Representative confocal images of WT and SynDIG4 KO neurons showing recycled and total GluA1 signals. (F) Quantification of the ratio of integrated density for recycled GluA1 to total GluA1 signals in WT and SynDIG4 KO neurons. Data are presented as mean ± SEM, with statistical significance determined using unpaired t-test; n =15-20; *p < 0.05. Scale bar, 10 µm.

### Retention of GluA1/GluA2-containing AMPARs in Rab4-positive endosomes in SynDIG4 deficient neurons

The above results suggest that AMPAR endosomal recycling is impaired in SynDIG4 KO neurons, though the specific intracellular compartments that SynDIG4 is involved in remains unclear. To identify the impaired compartments of AMPAR trafficking in SynDIG4 KO neurons, GluA1 and GluA2 subcellular distribution was investigated with different endosomal markers (39) in WT and SynDIG4 KO neurons. The results showed that the distribution of GluA1 and GluA2 in EEA1-positive endosomes were similar between WT and SynDIG4 KO neurons (**Figure 3A, 3D, S3A, S3D**), suggesting that trafficking to EEA1-positive early endosomes is unaffected, consistent with the results from the antibody-feeding assay (**Figure 2B, 2C**).

**Figure 3.**
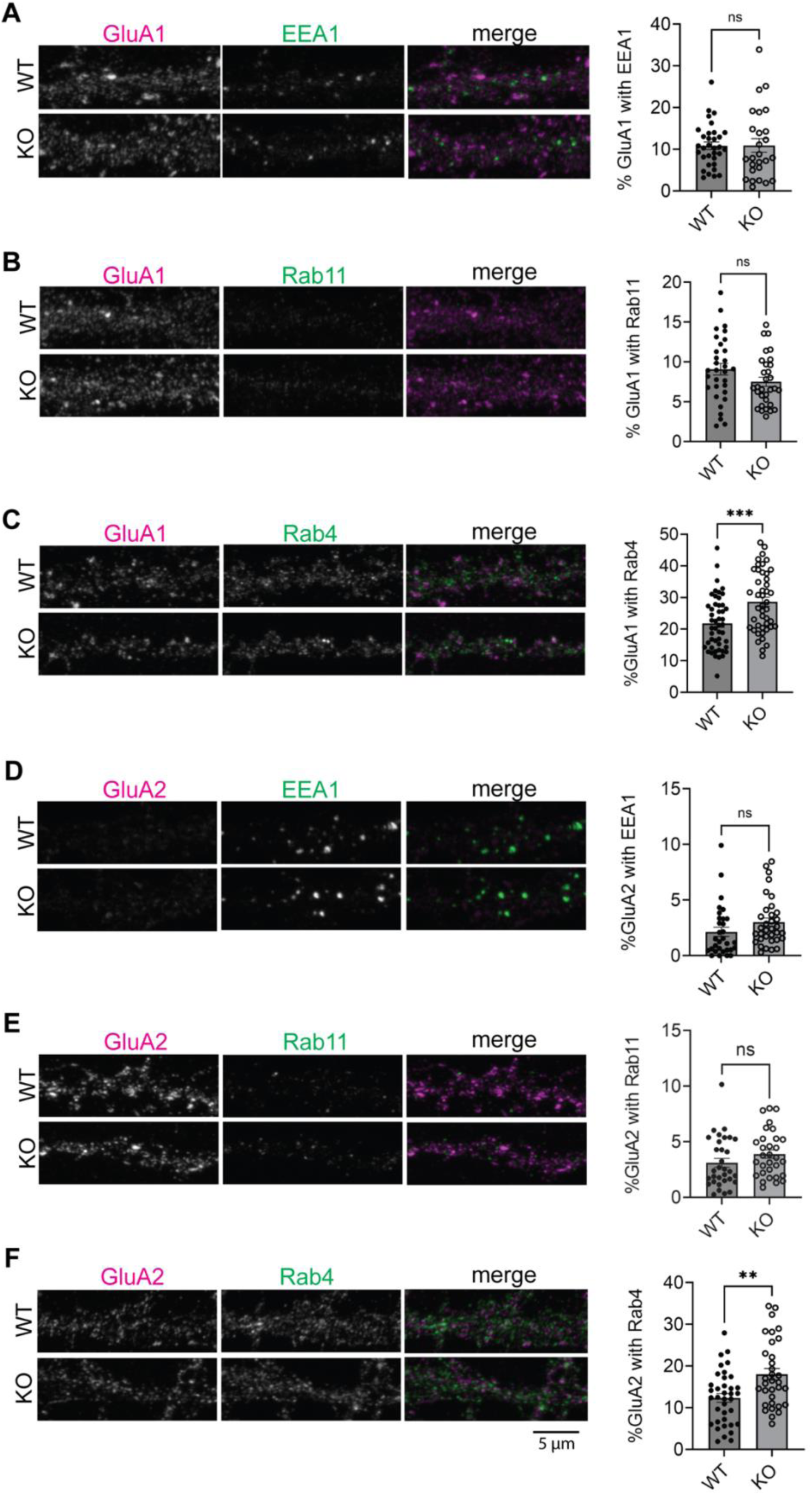
Accumulation of AMPARs in Rab4-positive endosomes in SynDIG4 KO neurons. (A) Representative images showing the colocalization of EEA1 and GluA1 and quantification of Manders’ colocalization coefficient for EEA1 and GluA1 in WT and SynDIG4 KO neurons. (B) Representative images showing the colocalization of Rab11 and GluA1 and quantification of Manders’ colocalization coefficient for Rab11 and GluA1 in WT and SynDIG4 KO neurons. (C) Representative images showing the colocalization of Rab4 and GluA1 and quantification of Manders’ colocalization coefficient for Rab4 and GluA1 in WT and SynDIG4 KO neurons. (D) Representative images showing the colocalization of EEA1 and GluA2 and quantification of Manders’ colocalization coefficient for EEA1 and GluA2 in WT and SynDIG4 KO neurons. (E) Representative images showing the colocalization of Rab11 and GluA2 and quantification of Manders’ colocalization coefficient for Rab11 and GluA2 in WT and SynDIG4 KO neurons. (F) Representative images showing the colocalization of Rab4 and GluA2 and quantification of Manders’ colocalization coefficient for Rab4 and GluA2 in WT and SynDIG4 KO neurons. Data are presented as mean ± SEM, with statistical significance determined using unpaired t-test; n = ∼40 dendrite stretches, 2–3 dendritic stretches were cropped from individual neuron images; **p < 0.01, ***p < 0.001. Scale bar, 5 µm.

AMPARs can be recycled to the plasma membrane through different endosomal trafficking pathways (39). To investigate which specific recycling process that SynDIG4 is involved in, we stained Rab4- and Rab11-positive compartments, markers for fast and slow-recycling endosomes, respectively (39), along with GluA1 or GluA2 in WT and SynDIG4 KO neurons. The results revealed that the levels of GluA1 and GluA2 in Rab11-positive endosomes were comparable between WT and SynDIG4 KO neurons (**Figure 3B, 3E, S3B, S3E**). In contrast, both GluA1 and GluA2 were significantly elevated in Rab4-positive endosomes in SynDIG4 KO neurons (**Figure 3C, 3F, S3C, S3F**). This result suggests that GluA1 and GluA2 are retained in Rab4-positive endosomes, preventing their transport back to the plasma membrane.

### Loss of SynDIG4 disrupts endosomal trafficking between Rab4- and Rab11-postive endosomes

Previous work proposed a model that impaired fusion between Rab4-positive and Rab11-positive endosomes leads to increased overlap between Rab4 and Rab11 compartments, thereby disrupting AMPAR trafficking (19). This disruption leads to a decrease in the surface expression of AMPARs and an increased colocalization of AMPARs with syntaxin13, a recycling endosome marker (20). To determine where SynDIG4 functions in this process, we co-stained for Rab4 and Rab11 with SynDIG4 in WT neurons. We found that the levels of SynDIG4 are comparable between Rab4- and Rab11-positive endosomes (**Figure 4A, S4A**), indicating that SynDIG4 localizes to both compartments. This finding suggests that SynDIG4 has functional roles at or between Rab4- and Rab11-positive endosomes. To test whether the trafficking between Rab4-positive and Rab11-positive endosomes is affected in SynDIG4 KO neurons, Rab4 and Rab11 were co-stained in WT and SynDIG4 KO neurons. Interestingly, the overlap between Rab4 signals and Rab11 signals was increased in SynDIG4 KO neurons, suggesting that endosomal trafficking between Rab4-positive and Rab11-postive endosomes is disrupted in SynDIG4 KO neurons (**Figure 4B, S4B**). At the same time, the overlap between EEA1 and Rab4 remained unchanged between WT and SynDIG4 KO neurons (**Figure 4C, S4C**). These results strengthen the idea that SynDIG4 functions in endosome recycling between Rab4-positive and Rab11-positive endosomes to regulate the distribution of AMPARs.

**Figure 4.**
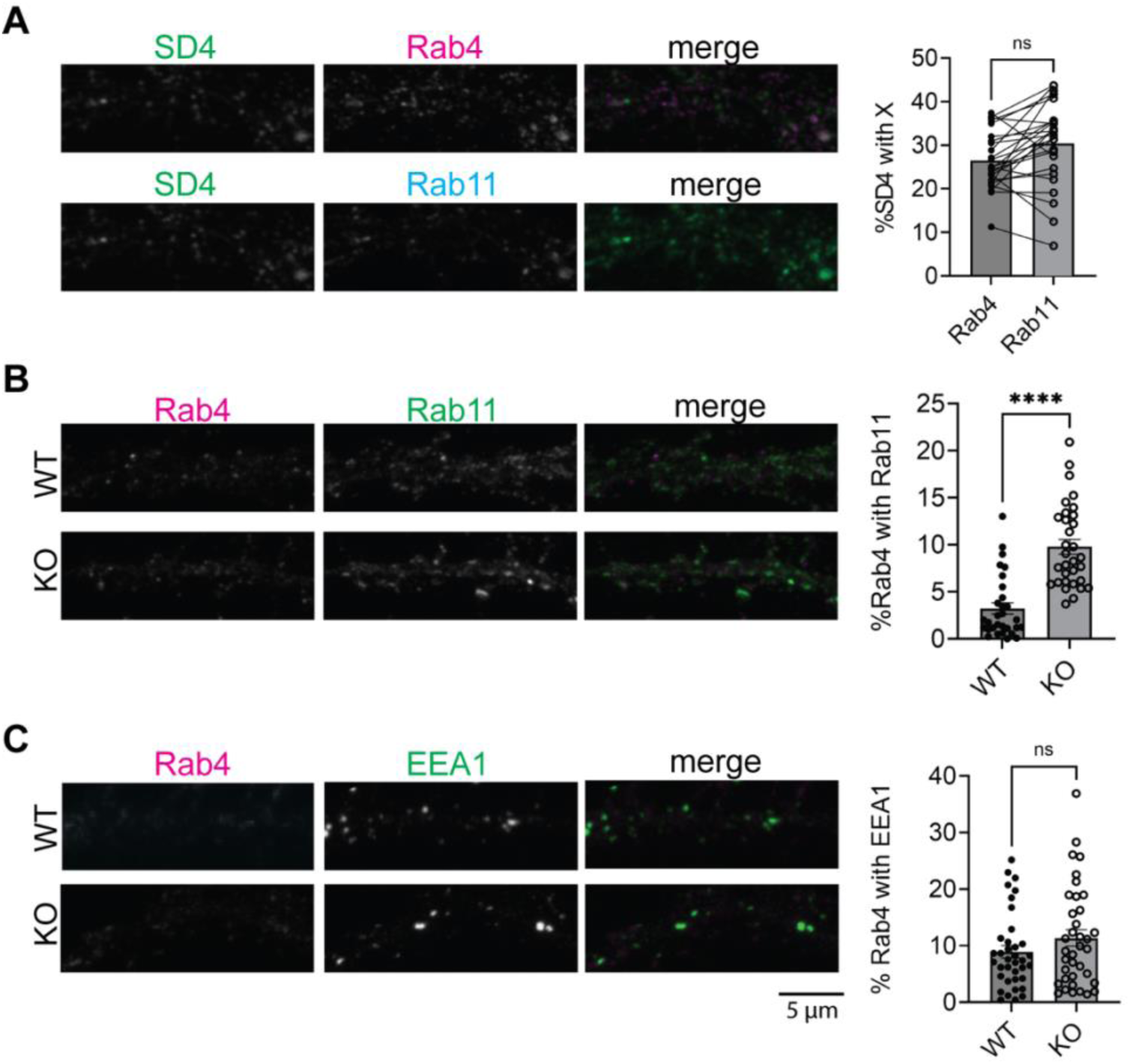
Increased colocalization of Rab4-positive and Rab11-positive vesicles in SynDIG4 KO neurons. (A) Representative images showing the colocalization of SynDIG4 and Rab4 or Rab11 and quantification of Manders’ colocalization coefficient for SynDIG4 and Rab4 or Rab11 in WT neurons. (B) Representative images showing the colocalization of Rab4 and Rab11 and quantification of Manders’ colocalization coefficient for Rab4 and Rab11 in WT and SynDIG4 KO neurons. (C) Representative images showing the colocalization of Rab4 and EEA1 and quantification of Manders’ colocalization coefficient for Rab4 and EEA1 in WT and SynDIG4 KO neurons. Data are presented as mean ± SEM, with statistical significance determined using unpaired t-test; n = ∼40 dendrite stretches, 2–3 dendritic stretches were cropped from individual neuron images; ****p < 0.0001. Scale bar, 5 µm.

### Decreased surface expression of GluA1-containing AMPARs at synaptic regions in SynDIG4 deficient neurons

We have demonstrated impaired AMPAR recycling in SynDIG4 KO neurons. Previous work showed decreased surface expression of GluA1 and GluA2 (32) and reduced density of extrasynaptic GluA1 and GluA2 puncta in SynDIG4 KO neurons (31). These findings suggest that SynDIG4 plays a critical role in both the surface expression and synaptic localization of AMPARs. To investigate SynDIG4 effects on the synaptic localization of surface AMPARs at the same time, sGluA1 was visualized using Stimulated Emission Depletion (STED) super-resolution microscopy to enhance resolution of molecular distributions within synapses by applying antibodies against GluA1-N to neurons prior to permeabilization (**Figure 5A**). As a control, antibodies against MAP2 added before permeabilization were undetectable, confirming that this surface-labeling protocol exclusively detects proteins located on the plasma membrane (**Figure S5**). The percentage of sGluA1 puncta overlapped with PSD95 puncta was decreased in SynDIG4 KO neurons (**Figure 5B**). To further investigate whether the association of sGluA1 with PSD95 was affected in SynDIG4 KO neurons, cross-correlation confidence analysis was performed. These results showed that the overlap of sGluA1 with PSD95 was reduced in SynDIG4 KO neurons (**Figure 5C**). Together, these results suggest that SynDIG4 is crucial for the synaptic localization of surface GluA1-containing AMPARs.

**Figure 5.**
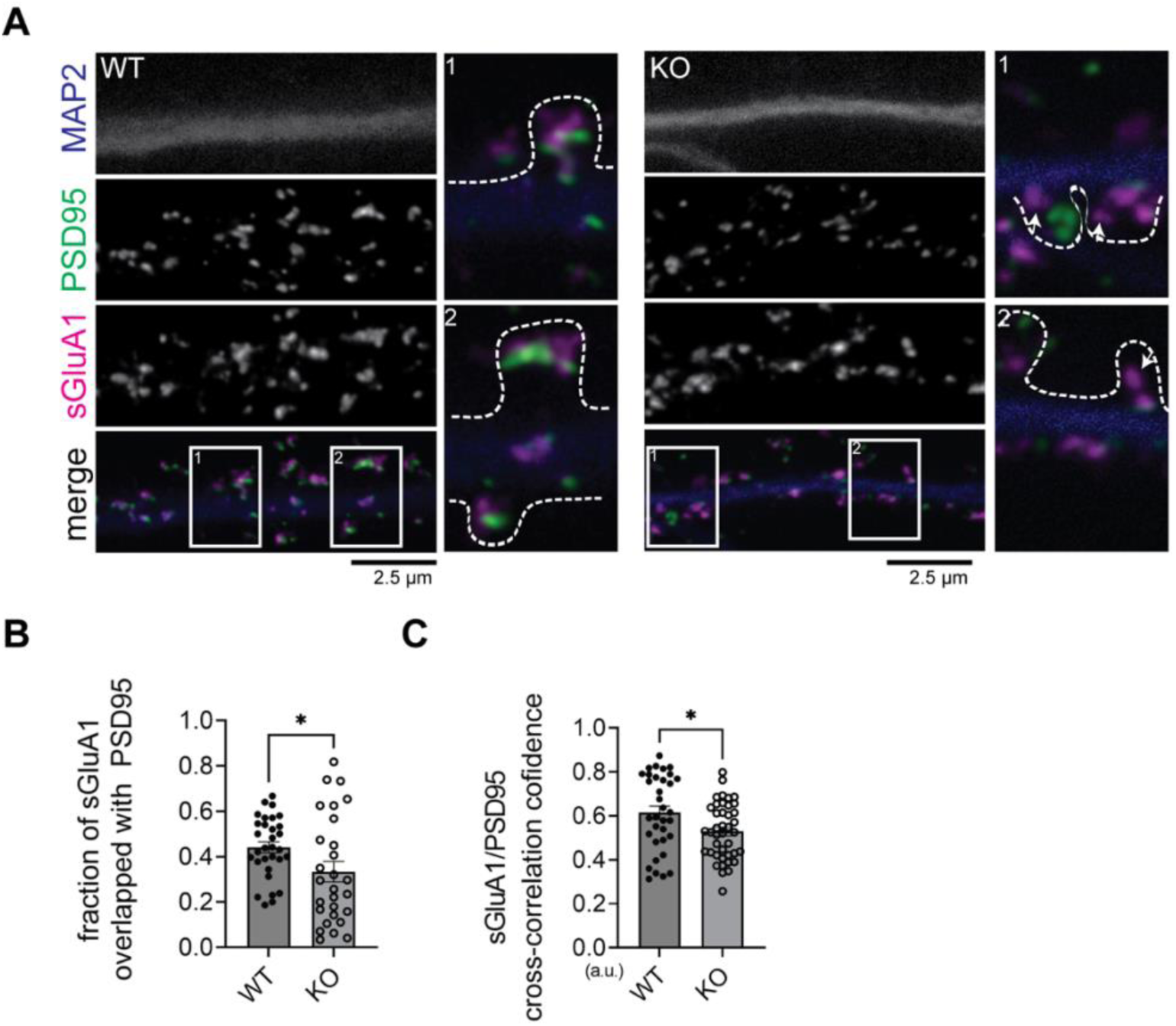
Association between sGluA1 and PSD95 was decreased in SynDIG4 KO neurons. (A) Representative STED images showing that sGluA1 is less associated or overlapped with PSD95 in SynDIG4 KO neurons (indicated by arrowhead). The outline of the spine is identified by the signals from the cell tracer DiA. (B, C) Quantitative analysis of sGluA1 and PSD95 colocalization using puncta overlap (B) and cross-correlation coefficient methods (C). Data are presented as mean ± SEM, with statistical significance determined using unpaired t-test; n = ∼40 dendrite stretches, 2–3 dendritic stretches were cropped from individual neuron images; *p < 0.05. Scale bar, 2.5 µm.

## DISCUSSION

Overall, our findings suggest that SynDIG4 promotes the surface expression of GluA1-containing AMPARs via the Rab4-dependent recycling pathway and regulates the synaptic localization of sGluA1. We found that the levels of endocytosed GluA1-containing AMPARs and their localization in early endosomes were comparable between WT and SynDIG4 KO neurons, indicating that SynDIG4 does not play a role in the endocytosis of GluA1-containing AMPARs under basal conditions. In contrast, recycling of GluA1-containing AMPARs was impaired in SynDIG4 KO neurons likely due to trafficking defects between Rab4- and Rab11-positive recycling endosomes, thereby reducing the levels of sGluA1 at synaptic sites (**Figure 6**).

**Figure 6.**
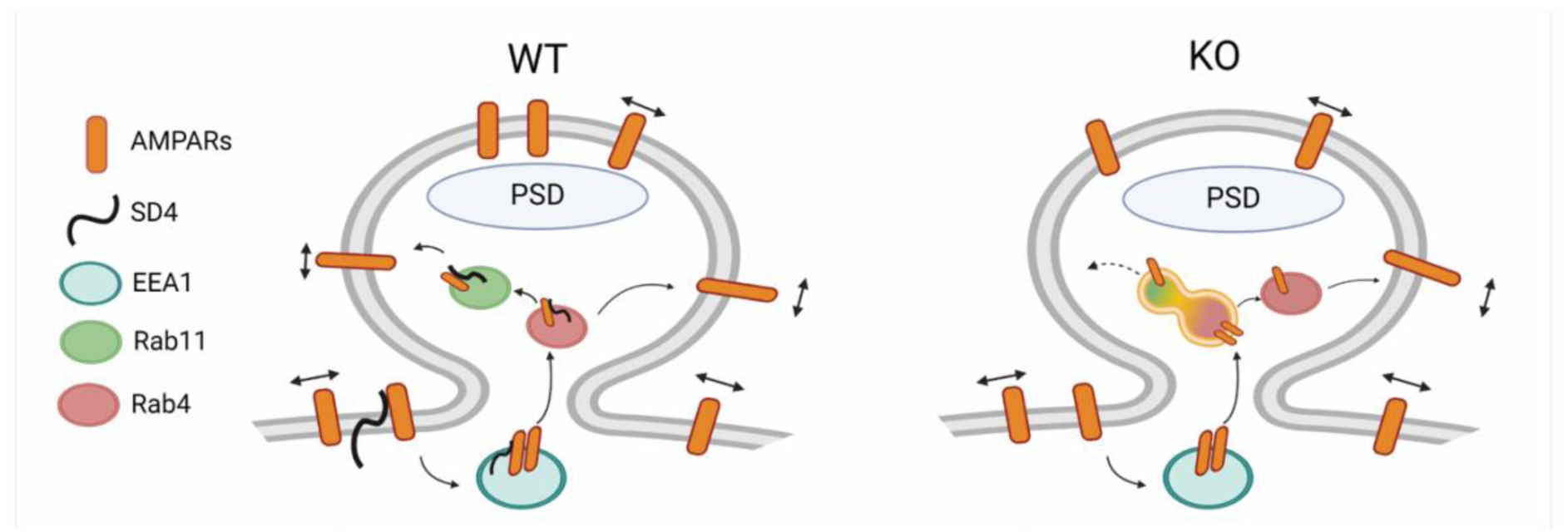
Model for the role of SynDIG4 in AMPAR distribution via endosomal recycling mechanism. In WT neurons, SynDIG4 (SD4) facilitates the trafficking of AMPARs between Rab4-positive and Rab11-positive endosomes under baseline conditions. In SynDIG4 KO neurons, the recycling of GluA1-containing AMPARs is reduced, and Rab11-positive endosomes fail to undergo fission from Rab4-positive endosomes. Consequently, the levels of intracellular AMPARs are increased, impairing the Rab4-Rab11-dependent recycling process. This SD4-mediated mechanism is crucial for the synaptic distribution of surface AMPARs. Created in BioRender. He, C. (2025) https://BioRender.com/k52o982.

### SynDIG4 promotes endosomal trafficking of AMPARs

Previous studies have shown that SynDIG4 co-localizes with the early endosomal marker EEA1 and with the transferrin receptor which is recycled between endosomes and the plasma membrane (34), suggesting that SynDIG4 is involved in recycling of AMPARs via the endosomal pathway. The endosomal pathway represents a collage of distinct yet overlapping compartments that are regulated by Rab proteins. Here we discovered that some SynDIG4 also overlaps with both the rapid and slow recycling endosomal markers Rab4 and Rab11, respectively (16). Rab11 is involved in the continuous recycling of endocytosed GluA1-containing AMPARs to the postsynaptic membrane via an EEA1-Rab4-Rab11 endosomal pathway (40–43); however, AMPARs can return to the surface directly from Rab4-positive compartments as was shown for GluA2 (17). Previous work proposed a model that impaired fusion between Rab4-positive and Rab11-positive endosomes leads to increased overlap between Rab4 and Rab11 compartments, thereby disrupting AMPAR trafficking (19). We demonstrate that upon loss of SynDIG4, the overlap between Rab4 and Rab11 increases significantly, which leads to a significantly increased accumulation of GluA1 and GluA2 in Rab4-positive recycling endosomes but not in Rab11-positive recycling endosomes, suggesting that SynDIG4 is involved specifically in the continuous and rapid recycling mediated by Rab4-dependent transport.

An interesting possibility is that SynDIG4 may be involved in mediating trafficking between Rab4 and Rab11 endosomes and point to a potential role for SynDIG4 in promoting the fission of Rab11-positive endosomes from Rab4-positive compartments. SynDIG4 belongs to a larger superfamily named “Dispanins” based on the prediction of two hydrophobic helical segments across the plasma membrane; however, later studies identified them as single pass type II transmembrane proteins with intracellular NT and extracellular CT, including SynDIG4 (33,34). Members of the Dispanin superfamily are thought to function as “fusogens”, facilitating or inhibiting membrane fusion (44). For example, interferon-induced transmembrane protein 3 (IFITM3), another member of the Dispanin superfamily, restricts viral entry in endosomes, by inhibiting fusion pore formation (45). Similarly, PRRT2 (Proline-Rich Transmembrane protein 2), to which SynDIG4 is related, has been shown to regulate the fusion of synaptic vesicles with the presynaptic plasma membrane, thereby controlling neurotransmitter release (46). Together, these findings strengthen the hypothesis that SynDIG4 contributes to trafficking between Rab4-positive and Rab11-positive endosomes by regulating the vesicle fusion and/or fission processes.

The retention of GluA1 and GluA2 in Rab4-positive endosomes, combined with possible defects in Rab4-Rab11 fission observed in SynDIG4 KO neurons, suggests that impaired endosomal trafficking between Rab4 and Rab11 disrupts the recycling of AMPARs. Interestingly, recycling of AMPARs through Rab4-Rab11-associated recycling endosomes and directly from Rab4-positive endosomes are both important for the supply for synaptic AMPARs (17,18). Moreover, the phosphorylation level of GluA1 at Serine 845 (S845), a site critical for AMPAR trafficking, was reduced in brain lysates from SynDIG4 KO mice (32). Consistent with this observation, SynDIG4 interacts with Protein Phosphatase 2B (PP2B, also known as calcineurin) (34), which is a phosphatase for GluA1 phosphorylation. Thus, SynDIG4 may regulate AMPAR trafficking through the interaction with PP2B. Our results support a model that SynDIG4 plays a pivotal role in AMPAR trafficking within the Rab4-Rab11 recycling pathway (**Figure 6**). However, we cannot exclude the possibility that SynDIG4 also contributes to AMPAR recycling via direct transport from Rab4-positive endosomes or through other potential mechanisms. For example, the increased GluA1 in Rab4-positive endosomes in SynDIG4 KO neurons could be due to a shift in trafficking pathways. That is, in response to the block in Rab4-Rab11-plasma membrane trafficking pathway, AMPAR transport moves to utilize the direct Rab4-plasma membrane pathway. neurons, which shifts the transport to the Rab4-direct pathway.

It is also possible that SynDIG4 regulates the trafficking of GluA1-containing AMPARs differently than GluA2-containing AMPARs, which may complicate interpretations of our results. SynDIG4 associates with both GluA1- and GluA2-containing AMPARs (26–28) and loss of SynDIG4 leads to decreased surface expression of GluA1 and GluA2 in brain lysates (32) and reduced density of extrasynaptic GluA1 and GluA2 in cultured neurons (31). Indeed, there are some aspects of SynDIG4 function that are selective for GluA1. In *Xenopus* oocytes, SynDIG4 was shown to modulate AMPAR gating properties in a subunit-dependent manner (31). Like other AMPAR auxiliary factors (21), SynDIG4 slows deactivation kinetics of both GluA1 homomers and GluA1/GluA2 heteromers. In contrast, SynDIG4 reduces desensitization only of GluA1 homomers and has no significant effect on desensitization of heteromeric GluA1/GluA2 (31). In native cryo-EM structures, SynDIG4 association with AMPARs appears to be through the M4 helix of GluA1 (29). Intriguingly, both NMDAR-dependent LTD (32) and single tetanus-induced LTP (31), which require calcium-permeable AMPARs (35), which are mostly GluA1 homomers, are impaired in SynDIG4 deficient neurons while baseline synaptic transmission is intact. Thus, it is possible that SynDIG4 is required for trafficking of both GluA1/GluA2 heteromers as well as GluA2 homomers at baseline through independent endosomal pathways. Additional experiments beyond the scope of this study are necessary to investigate this interesting possibility.

### SynDIG4 regulates the distribution of surface AMPARs at synapses

Using STED microscopy, we found that the overlap between sGluA1/sGluA2 and PSD95 were decreased in SynDIG4 KO neurons. These findings suggest a reduction in sGluA1/sGluA2-containing AMPARs at or near PSD95, consistent with the observed decrease in miniature excitatory postsynaptic current (mEPSC) amplitude in SynDIG4 KO mice (31). Interestingly, under confocal microscopy, the density of GluA1 puncta was increased in synaptic regions identified by overlap with the presynaptic marker vGLUT1 (31). These findings indicate that while total GluA1 level at synapses is increased, its surface expression was reduced, implying that inGluA1 may be retained within the synaptic region. However, this interpretation should be considered with caution, as different imaging methods (STED vs. confocal) and distinct synaptic markers (PSD95 vs. vGLUT1) were used, which could lead to potential misinterpretations.

### Role of SynDIG4 in endocytosis of AMPARs

Interestingly, our prior study investigating the distribution of SynDIG4 and AMPARs in heterologous cells proposed that endocytosis is likely required for the clustering of live-labeled GluA1 and GluA2, but not GluK2, with SynDIG4 (30). This raises the possibility that SynDIG4 may be involved in the endocytosis-dependent clustering of GluA1/GluA2 on the plasma membrane, despite not affecting the overall amount of AMPARs being internalized. Alternatively, SynDIG4 and AMPARs may undergo endocytosis independently and associate within endosomes to promote bi-directional clustering. Importantly, the clustering experiments were conducted at 37°C, a temperature that facilitates both endocytosis and exocytosis, leaving open the possibility that exocytosis also contributes to GluA1/GluA2 clustering in heterologous cells. Our findings here demonstrate that SynDIG4 is dispensable for the internalization of GluA1-containing AMPARs at baseline. However, they do not exclude the possibility that SynDIG4 influences AMPAR trafficking through mechanisms involving endocytosis such as during synaptic plasticity.

In summary, our findings demonstrate that SynDIG4 plays a critical role in the endosomal trafficking of GluA1-containing AMPARs via Rab4-positive recycling pathways and is essential for the proper distribution of surface AMPARs at synapses. Furthermore, SynDIG4 KO mice display deficits in LTD (32) and single-tetanus LTP (31), both of which require GluA1. SynDIG4 KO mice also display deficits in hippocampal-dependent learning and memory (31), and downregulation of SynDIG4 has been found in patients with Alzheimer’s disease (47). These findings highlight the importance of investigating SynDIG4 function in synaptic plasticity, its contributions to learning and memory, and its potential role in the pathology of Alzheimer’s disease. These areas are the focus of our current research.

## DATA AVAILABILITY STATEMENT

The raw data supporting the conclusions of this article will be made available by the authors, without undue reservation.

## ETHICS STATEMENT

The animal study was approved by Institutional Animal Care and Use Committee of the University of California, Davis. The study was conducted in accordance with the local legislation and institutional requirements.

## AUTHOR CONTRIBUTIONS

C-WH and ED: methodology and writing. C-WH: investigation. C-WH and ED: formal analysis. ED: funding acquisition and supervision. Both authors reviewed the article and approved the final submitted version.

## ACKNOWLEDGEMENTS

We extend our sincere gratitude to Drs. Johannes Hell, John Gray, and Madeline Nieves-Cintrón (University of California, Davis) for their invaluable insights and feedback throughout the research process. We thank members of the Díaz Lab, particularly Dr. David Speca, for their constructive suggestions on the research and comments on the manuscript. We are especially grateful to Dr. Ingrid Brust-Mascher (Advanced Imaging Facility, University of California, Davis School of Veterinary Medicine) for her expert assistance with fluorescent microscopy. Additionally, we thank Dr. James Trimmer and members of Dr Hell’s lab, with special recognition to Zoila Estrada-Tobar, (University of California, Davis), for generously sharing antibodies and reagents used in this research.

