## Supplemental Figures for "Loss of SynDIG4/PRRT1 alters distribution of AMPA receptors in Rab4- and Rab11-positive endosomes and impairs basal AMPA receptor recycling"

*Supplementary Material*

### Supplementary Figures

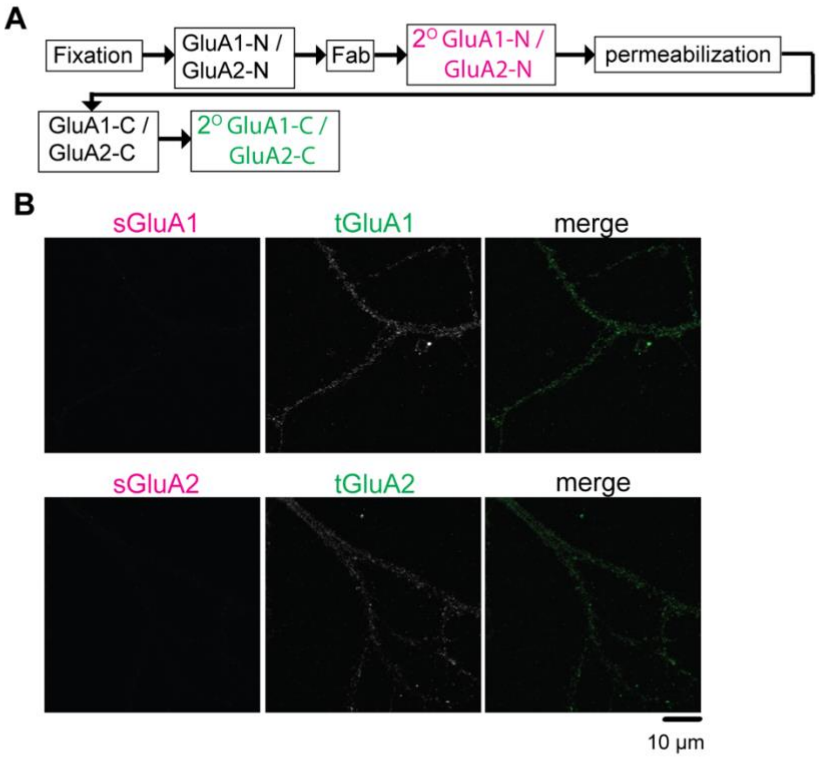


**Figure S1. Fab treatment blocks binding sites for anti-mouse IgG antibodies**
(A) Schematic diagram illustrating the experimental procedure for Fab treatment. After GluA1-N or GluA2-N primary antibody incubation, Fab was applied to the neurons to prevent binding of fluorescently conjugated secondary anti-mouse IgG antibodies. (B) Representative confocal images demonstrating that surface GluA1 (sGluA1) and surface GluA2 (sGluA2) signals are undetectable following Fab application. In contrast, total GluA1 (tGluA1) and total GluA2 (tGluA2) signals are detectable upon permeabilization and labeling with primary GluA1-C or GluA2-C antibodies and appropriate secondary antibodies. Scale bar, 10 µm.

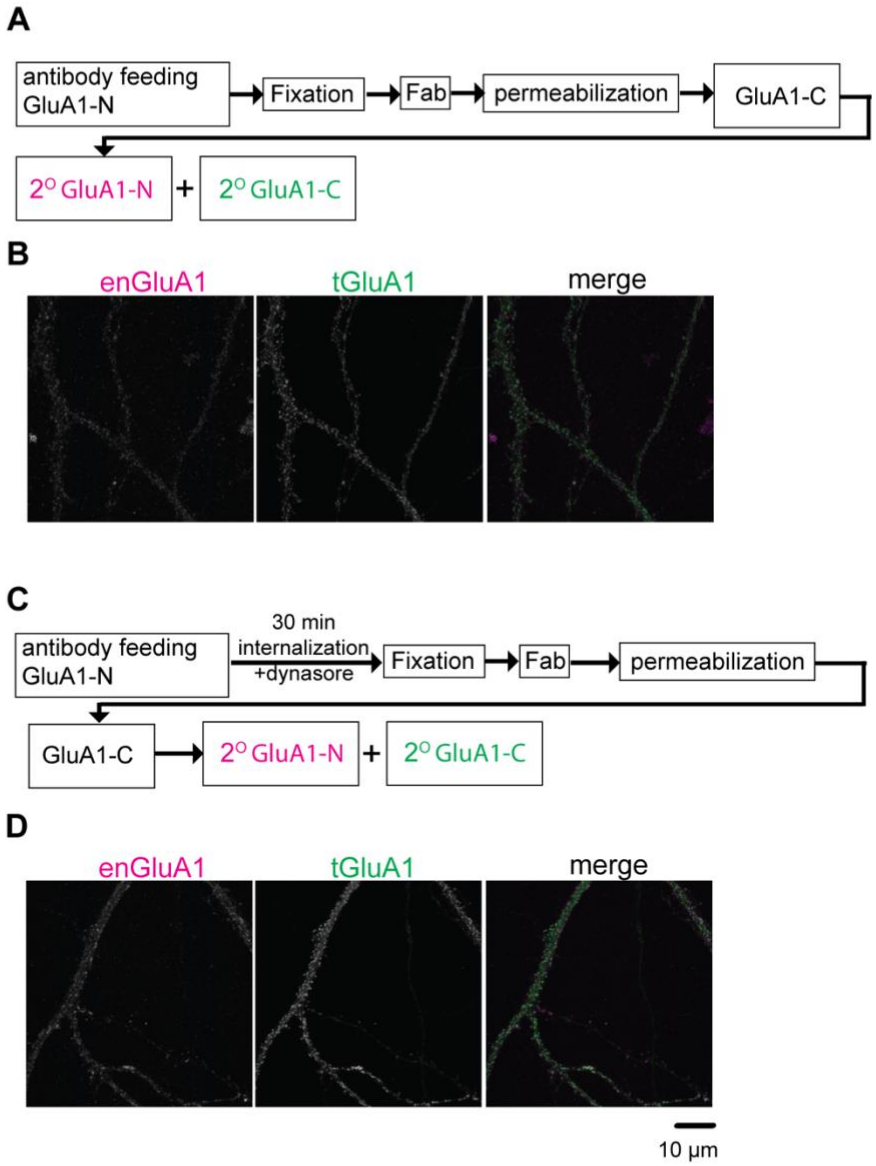


**Figure S2. Dynasore inhibits dynamin-dependent endocytosis of GluA1**

(A) Schematic diagram illustrating the experimental procedure. After antibody feeding (GluA1-N) at room temperature for 20 min, neurons were fixed and then incubated with Fab antibodies to block epitopes on the cell surface. (B) Minimal endocytosis of GluA1-N-labeled AMPARs (enGluA1) occurred during the antibody-feeding process. Total GluA1 (tGluA1) was immunostained for comparison. (C) Schematic diagram showing the experimental procedure where dynasore was applied during the 30 min internalization period. (D) Representative images show a reduction in endocytosed GluA1 (enGluA1) signals following dynasore treatment, suggesting inhibition of the dynamin-dependent endocytosis process. Total GluA1 (tGluA1) was immunostained for comparison. Scale bar, 10 μm.

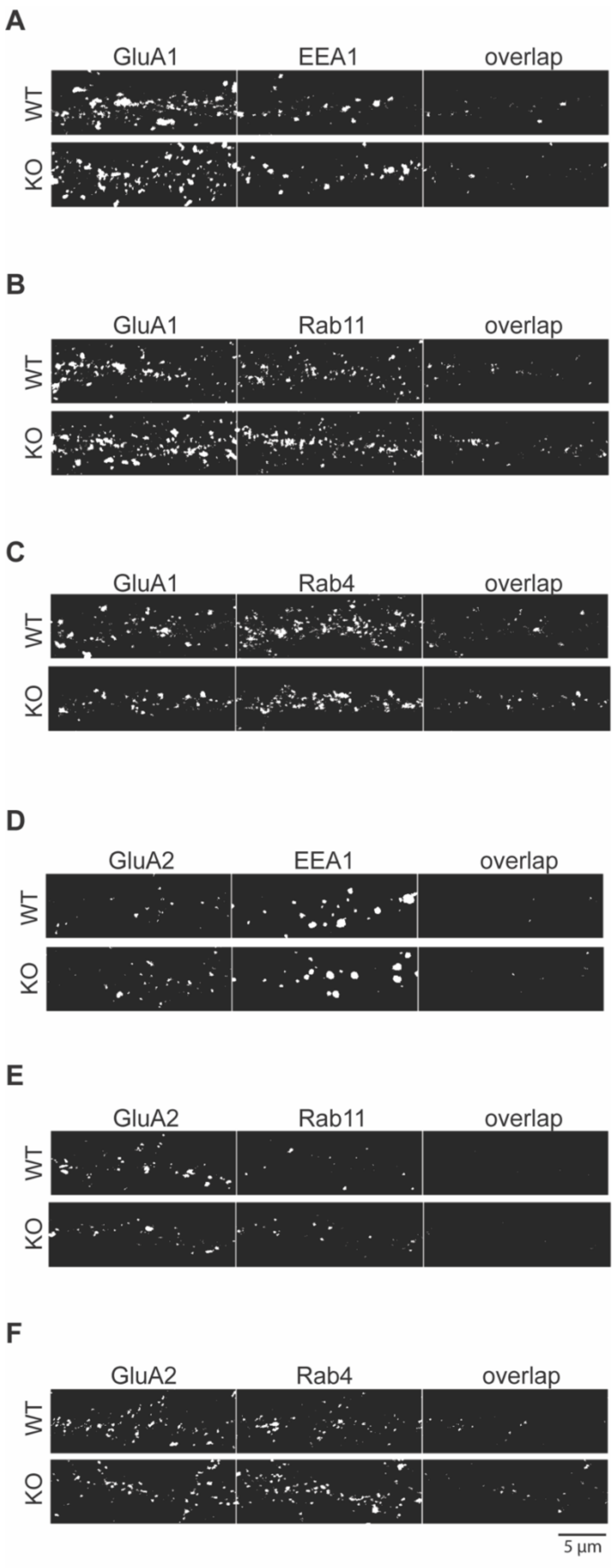
A
GluA1
EEA1
overlap
WT
KO
B
GluA1
Rab11
overlap
WT
KO
C
GluA1
Rab4
overlap
WT
KO
D
GluA2
EEA1
overlap
WT
KO
E
GluA2
Rab11
overlap
WT
KO
F
GluA2
Rab4
overlap
WT
KO
5 µm

**Figure S3. Overlap between GluA1 or GluA2 and endosomal markers.**
(A-C) Representative thresholded images showing the overlap of GluA1 and EEA1 (A), Rab11 (B), or Rab4 (C) in wild-type (WT) and SynDIG4 knockout (KO) neurons used for analysis in Figure 3A-C. (D-F) Representative thresholded images showing the overlap of GluA2 and EEA1 (D), Rab11 (E), or Rab4 (F) in WT and SynDIG4 KO neurons used for analysis in Figure 3D-F. Scale bar, 5 μm.

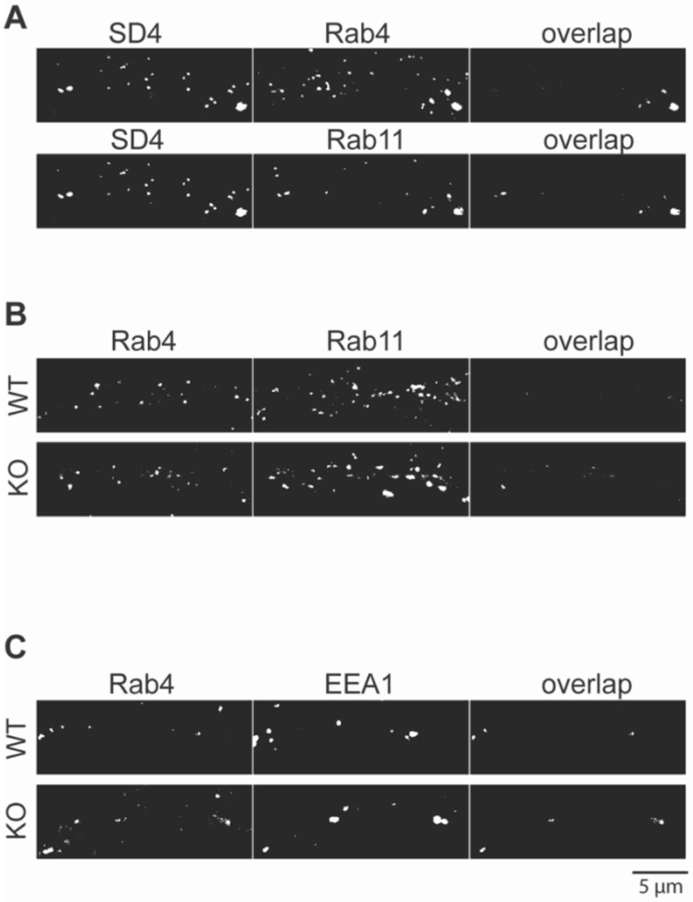


**Figure S4. Overlap between SynDIG4 and endosomal markers.**
(A) Representative thresholded images showing the overlap of SynDIG4 and Rab4 or Rab11 in wild-type (WT) neurons used for analysis in Figure 4A. (B) Representative thresholded images showing the overlap of Rab4 and Rab11 in WT and SynDIG4 knockout (KO) neurons used for analysis in Figure 4B. (C) Representative thresholded images showing the overlap of Rab4 and EEA1 in WT and SynDIG4 KO neurons used for analysis in Figure 4C. Scale bar, 5 μm.

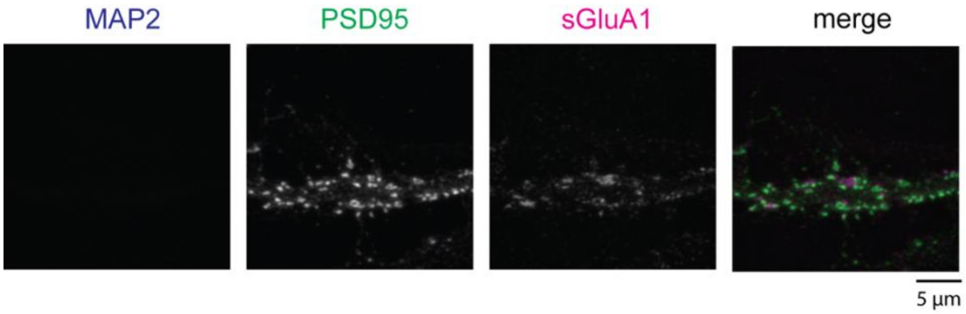


**Figure S5. Surface labeling control with anti-MAP2 antibody**.
Antibodies against GluA1-N and MAP2 were applied to neurons prior to permeabilization. Neurons were fixed and permeabilized followed by immunostaining with antibodies against PSD95, an intracellular synaptic scaffold, and subsequent fluorescently conjugated secondary antibodies for GluA1-N, MAP2, and PSD95. Representative confocal images demonstrate that the anti-MAP2 antibody does not penetrate the plasma membrane prior to permeabilization, confirming that the surface-labeling protocol specifically detects proteins localized on the plasma membrane, while excluding intracellular epitopes. Scale bar, 5 µm.
